# Estimates of persistent inward currents are reduced in upper limb motor units of older adults

**DOI:** 10.1101/2021.06.18.448899

**Authors:** Altamash S Hassan, Melissa E Fajardo, Mark Cummings, Laura Miller McPherson, Francesco Negro, Julius P A Dewald, C J Heckman, Gregory E P Pearcey

**Author notes:** <u>Address for correspondence</u>: Dr. Gregory Pearcey, Department of Physical Therapy & Human Movement Sciences, 645 N. Michigan Ave., Suite 1100, Northwestern University, Chicago, IL 60611.

## Abstract

Aging is a natural process that causes alterations in the neuromuscular system, which contribute to weakness and reduced quality of life. Reduced firing rates of individual motor units (MUs) likely contribute to weakness, but the mechanisms underlying reduced firing rates are not clear. Persistent inward currents (PICs) are crucial for the initiation, gain control, and maintenance of motoneuron firing, and are directly proportional to the level of monoaminergic input. Since the concentration of monoamines (i.e. serotonin and norepinephrine) are reduced with age, we sought to determine if estimates of PICs are reduced in older (>60 years old) compared to younger adults (<35 years old). We decomposed MU spike trains from high-density surface electromyography over the biceps brachii and triceps brachii during isometric ramp contractions to 20% of maximum. Estimates of PICs (i.e. ΔF) were computed using the paired MU analysis technique. Regardless of the muscle, peak firing rates of older adults were reduced by ~1.6 pulses per second (pps) (*P = 0.0292*), and ΔF was reduced by ~1.9 pps (*P < 0.0001*), compared to younger adults. We further found that age predicted ΔF in older adults (*P = 0.0261*), resulting in a reduction of ~1pps per decade, but there was no relationship in younger adults (*P = 0.9637*). These findings suggest that PICs are reduced in older adults, and, further, age is a significant predictor of estimates of PICs in older adults. Reduced PIC magnitude represents one plausible mechanism for reduced firing rates and weakness in older individuals.

**KEY POINTS:**

- Persistent inward currents play an important role in the neural control of human movement and are influenced by neuromodulation via monoamines originating in the brainstem.
- During aging, motor unit firing rates are reduced, and there is deterioration of brainstem nuclei, which may reduce persistent inward currents in alpha motoneurons.
- Here we show that estimates of persistent inward currents (ΔF) of both elbow flexor and extensor motor units are reduced in older adults.
- Estimates of persistent inward currents have a negative relationship with age in the older adults, but not young.
- This novel mechanism may play a role in alteration motor firing rates that occurs with aging, which may have consequences for motor control.

## INTRODUCTION

Aging is a natural process that causes alterations within the neuromuscular system, which can have severe consequences on health and quality of life in older adults (McNeil & Rice, 2018). Even in the absence of disease, there is age-related loss of muscle or lean mass (i.e. sarcopenia), and, perhaps more importantly, age-related loss of strength (i.e. dynapenia). Emerging evidence suggests that dynapenia is a significant contributor to quality of life in the elderly (Mitchell *et al.*, 2012). Indeed, biophysical properties of the muscle play a role in the reduced force generating capacity, but neural factors are likely to contribute as well.

There is a progressive loss in the number of motor units (MUs) with age (McNeil *et al.*, 2005), which comprise the muscle fibers and their parent motoneurons (Heckman & Enoka, 2012). As such, death of motoneurons is widely accepted as a precursor for many of the age-related adaptations in the nervous system (McNeil & Rice, 2018). Following death of motoneurons, the nervous system displays astounding plasticity, as evidenced by the reinnervation of orphaned muscle fibers by axonal sprouting (Gordon *et al.*, 2004), a process known as MU remodeling (Hepple & Rice, 2016). Since it is speculated that the loss of larger/faster motoneurons precedes the loss of smaller/slower type motoneurons (Kanda & Hashizume, 1989), reductions in MU firing rates are typically ascribed to this mechanism. Dalton et al. (Dalton *et al.*, 2010) previously showed that the firing rates of both biceps brachii (BIC) and triceps brachii (TRI) MUs are reduced across a wide range of contraction intensities in older, compared to young, adults. They suggested that the relatively higher proportional loss of higher threshold motoneurons may play a role in the age-related decline in firing rates, but alterations in the biophysical properties of the motoneurons are also likely to contribute.

Altered intrinsic motoneuron excitability may play a major role in age-related changes in motoneuron firing patterns. Although motoneurons were once believed to integrate their synaptic inputs passively, many studies have demonstrated that this integration is a highly active process due to voltage-sensitive ion channels in their dendrites (Heckman *et al.*, 2008a; Heckman *et al.*, 2008b). Persistent inward currents (PICs) amplify and prolong excitatory synaptic input to the motoneuron (Lee & Heckman, 1998, 2000), which are the result voltage-gated slow activating L-type Ca^2+^ and fast activating persistent Na^+^ currents (Heckman *et al.*, 2008b). PICs are activated near threshold and can amplify synaptic currents by as much as 3-5 fold (Binder & Powers, 2001), and the level of PIC activation is highly dependent on the neuromodulatory drive from the monoaminergic system (i.e. serotonergic and noradrenergic drive) (Lee & Heckman, 1998, 2000). In addition, PICs are reduced with antagonist muscle afferent input (i.e. reciprocal inhibition), illustrating a role for inhibition in the control of PIC activity (Heckman *et al.*, 2008a; Powers *et al.*, 2012). Therefore, changes in levels of monoaminergic drive, intrinsic motoneuron excitability (i.e. monoamine receptor or ion channel function), and inhibition may alter MU firing patterns, as well as estimates of PICs, with age (Johnson *et al.*, 2017).

Furthermore, recent work has called for the investigation of PIC estimates in the aging neuromuscular system (Latella, 2021). The function of two primary monoaminergic nuclei in the brainstem, the raphe nuclei and locus coeruleus, have been shown to deteriorate with age (Shibata *et al.*, 2006; Pagano *et al.*, 2017). This deterioration likely results in reduced monoaminergic drive, and consequently reduced activation of PICs.

Fortuitously, recent advances in technology have enabled us to sample from large populations of concurrently active MUs, by using high-density surface electromyography (HD-sEMG) array electrodes and blind source separation algorithms (Holobar & Zazula, 2007; Negro *et al.*, 2016) with great success (Yavuz *et al.*, 2015; Del Vecchio *et al.*, 2018; Thompson *et al.*, 2018; Cogliati *et al.*, 2020; Del Vecchio *et al.*, 2020; Hassan *et al.*, 2020; Kim *et al.*, 2020; Martinez-Valdes *et al.*, 2020). This non-invasive technology has created an opportunity to further study the age-related alterations in the neuromuscular system by sampling from many concurrently active MUs. Using this technology allows us to gain better appreciation of the population behaviour and provide more insights about the control of large portions of the motor pool, which was difficult to achieve with intramuscular EMG approaches.

In this study, we examined whether the MU firing characteristics in a large population of concurrently active MUs differed between a group of younger and older adults. More specifically, we compared MU firing patterns of the elbow flexor and extensors during triangular isometric contractions. We hypothesized that since previous work has shown MU firing rates are reduced in older adults, we would observe reductions in peak firing rates of both BIC and TRI MUs, as well as estimates of PIC magnitude (i.e. ∆F), in the older group. In addition, since healthy younger adults are unlikely to have an impairment in PIC function, we investigated the relationship between age and estimates of PICs in both groups separately. We hypothesized that there would be no relationship between age and estimates of PICs in the younger group, however there would be a negative relationship between age and estimates of PICs in the older group.

## METHODS

### Participants

In order to compare MU firing behaviour between healthy younger and older individuals, we recruited 10 younger (26 [2.87] years old, 3 female) and 10 older (67 [4.40] years old, 2 female) adults. At the time of testing, all participants were free of neurological, motor, and muscular impairments. All participants provided written informed consent in accordance with the Declaration of Helsinki, which was approved (STU00084502-CR0003) by the Institutional Review Board of Northwestern University.

### Experimental Apparatus

The experimental apparatus, protocol, and data processing methods utilized is similar to those used in a previous experiment in our lab (Hassan *et al.*, 2020). Participants were secured in a Biodex chair (Biodex Medical Systems, Shirley, NY) with their dominant upper limb rigidly fixed to a six degree-of-freedom load cell with a fiberglass cast (JR3, Inc., Woodland, CA). The casted arm was positioned at a shoulder abduction angle of 75° and an elbow flexion angle of 90°. An illustration of the experimental setup is shown in Figure 1.

**Figure 1:**
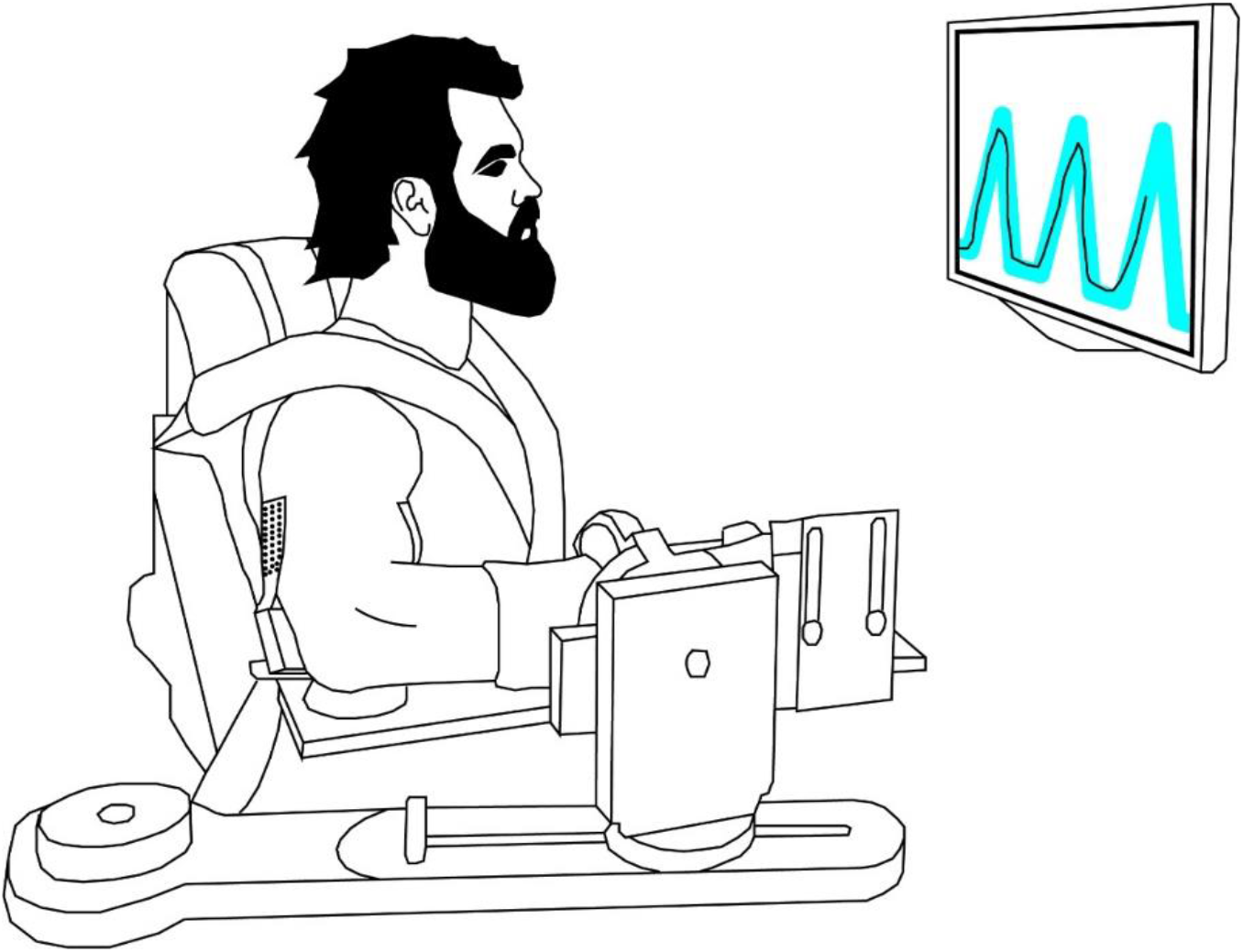
An illustration of the experimental setup. High-densit electromyography (HD-sEMG) arrays were placed on the latera the TRI and along the muscle belly of the BIC.

High-density surface EMG (HD-sEMG) arrays (64 electrodes, 13×5, 8mm I.E.D., GR08MM1305, OT Bioelettronica, Inc., Turin, IT) were placed on the BIC and the lateral head of the TRI on the casted limb. HD-sEMG data were sampled (2048 Hz), amplified (x150), and band-pass filtered (10-500Hz) using a Quattrocentro signal amplifier (OT Bioelettronica, Inc., Turin, IT). A reference electrode was placed on the acromion process of the casted arm. Prior to collecting experimental data, real-time HD-sEMG recordings were checked visually to ensure high signal-to-noise ratios.

Torque was sampled at 1024 Hz with a forearm-load cell interface. The limb segment lengths and joint angles were converted into elbow flexion and extension torques using a Jacobian based algorithm implemented by a custom MATLAB software (MathWorks). As EMG and force/torque recordings were collected using separate computers, a 1 second TTL pulse was transmitted to both computers for data alignment. Each trial was temporally synced offline using cross-correlation of the TTL pulses.

### Experimental Protocol

Participants were initially asked to produce their maximum voluntary torques (MVTs) in the directions of elbow flexion or extension. A wall-mounted computer monitor placed directly in front of participants provided real-time feedback of torque performance. MVT trials were repeated until three trials in which the peak torque was within 10% of each other were collected. If the last trial had the highest peak torque, an additional MVT trial was collected. Participants were verbally encouraged during MVT trials to ensure peak torque performance and were given adequate rest between trials to prevent muscle fatigue.

Each of the subsequent experimental trials consisted of three triangular isometric elbow extension/flexion torque ramps, separated by 10 seconds of rest. Each ramp required participants to increase torque (2% MVT/s) to 20% MVT over 10 seconds, and then decrease (−2% MVT/s) to 0% MVT over the next 10 seconds. During all trials, real-time torque feedback, as well as the desired experimental torque profile, were provided to the participants. In order to avoid fatigue, a minimum of 2 minutes of rest was given to participants between trials. Trials that did not exhibit a smooth increase of torque from 0% to 20% MVT and a smooth decrease of torque from 20% to 0% MVT over the desired timeframe were discarded, as were trials that display any sudden jerks in torque.

### Data Analysis

#### MU Decomposition & Variables of Interest

After data acquisition, each EMG channel of the surface array was manually inspected and any channels with substantial artifacts, noise, or analog to digital saturation were removed. A convolutive blind-source separation algorithm (Negro *et al.*, 2016) with a silhouette threshold of 0.85 was used to decompose HD-sEMG into individual MU spike trains. All decomposed MU spike trains were visually inspected for each participant and trial through a custom-made graphical user interface in MATLAB. Any minor errors were corrected by local re-optimization of decomposition parameters in a manner similar to recent studies using the same blind source separation algorithm (Boccia *et al.*, 2019; Afsharipour *et al.*, 2020; Del Vecchio *et al.*, 2020; Hassan *et al.*, 2020; Martinez-Valdes *et al.*, 2020). Instantaneous MU firing rates were calculated as the inverse of the interspike intervals of each MU spike train and smoothed using a 2 second Hanning window using a custom-written MATLAB script.

Peak, total duration, and total range of smoothened decomposed MU firing rates were extracted through custom written MATLAB scripts for each muscle. MU range and duration during the ascending and descending phases of torque production were also extracted (see Figure 2).

**Figure 2:**
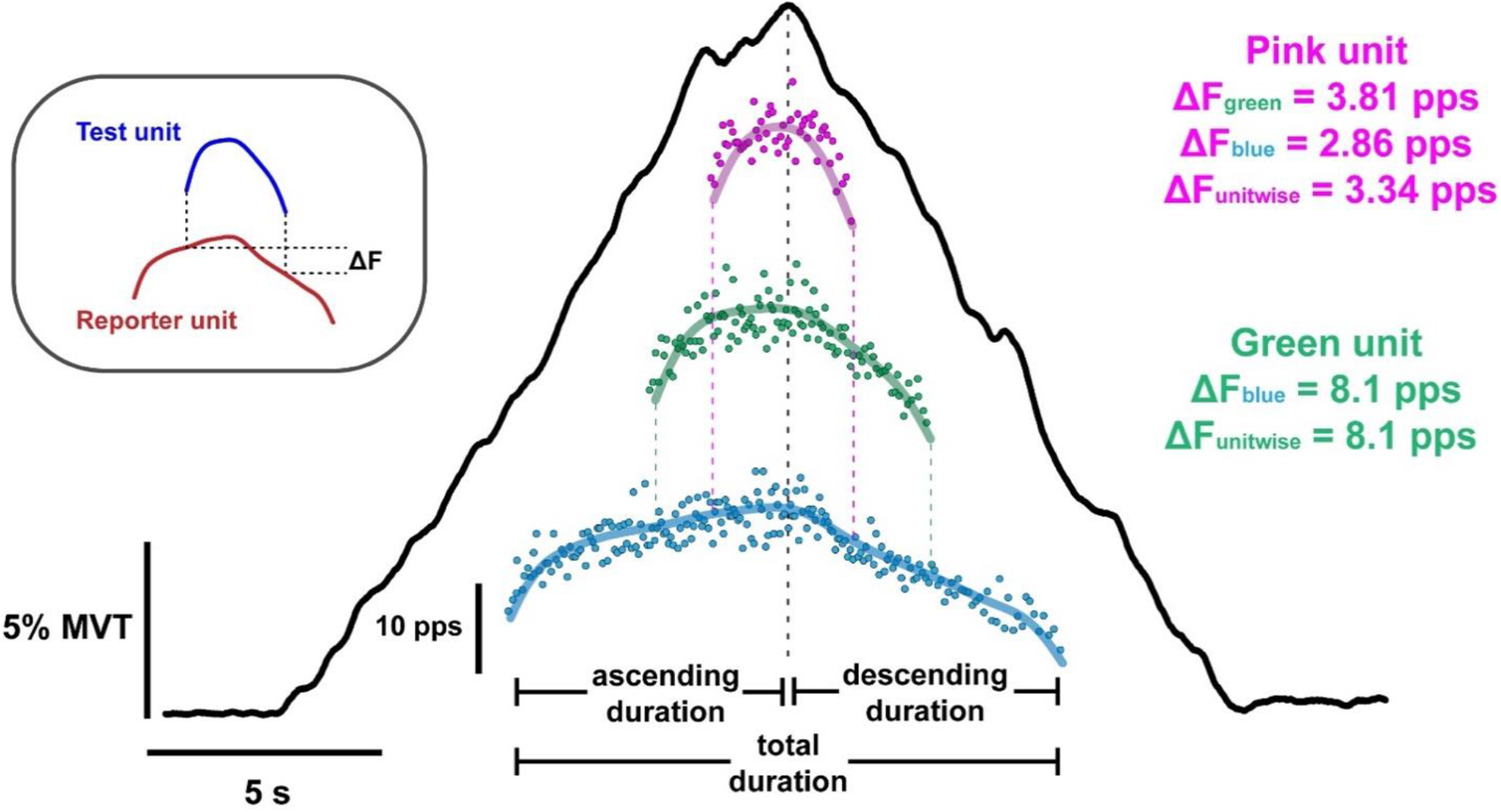
Top left shows the paired MU analysis method used to estimate persistent inward current magnitude which quantifies the onset-offset hysteresis (i.e. ΔF) of a higher threshold (test unit; blue) MU with respect to a lower threshold (reporter unit; red). In the center, a typical extension torque trace (black line) from one younger participant is shown. Underneath the torque ramp, 3 out of the 12 decomposed MU firing patterns are shown. Each point indicates the instantaneous firing rate for each interstimulus interval, and the thick coloured lines indicate the smoothed firing rates of each MU. At the ends of the green and pink unit are vertical dotted lines that extend downward to the units below them (i.e. recruited at lower torque), which helps indicate the onset and offset of firing with respect to their reporter units. Individual ΔF values obtained from each reporter unit, and the mean of those values (ΔF unitwise) is shown for each test unit. The vertical black dotted line indicates peak torque in this ramp for ease of viewing the time point between ascending and descending duration.

#### Estimating Persistent Inward Currents (PICs)

The effects of persistent inward currents (PICs) on motoneuronal firing patterns can be appreciated through MU onset-offset hysteresis. The best approximation of this hysteresis, Delta F (ΔF), is calculated as the difference in the smoothed firing rate of a reporter (lower threshold) MU between the times of recruitment/derecruitment of a test (higher threshold) MU (Gorassini *et al.*, 2002). All ΔF values used throughout this study are “unitwise” mean values. That is, the average of ΔF values obtained for each test unit from all possible reporter units (see Figure 2). Criteria for inclusion of ΔF values from MU pairs were that the test MU was 1) recruited at least 1 second after the control unit to ensure full activation of PIC, and 2) derecruited at least 1.5 seconds prior to the control MU to prevent ΔF overestimation (see Hassan et al. (Hassan *et al.*, 2020) for more details on utilizing ΔF for paired MU analysis).

### Statistical Analysis

All data was imported into GraphPad (version 9.0.1 for Windows, GraphPad Software, San Diego, California USA) where descriptive statistical analyses were performed. Hedge’s g effect sizes (ES) were calculated to provide a standardized effect for the mean differences between the younger and older subjects for each variable. Mean and standard deviation values for each variable reported are group means, which represent the average and error of the individual means computed for each participant.

We detail the effects of healthy aging on MU firing characteristics using linear mixed effects models. More specifically, we take into consideration all of our data points rather than averaging across them and basing our analysis on the mean within an individual trial or subject (Giboin *et al.*, 2020). All of these analysis were performed in R (R Core Team 2020, R Foundation for Statistical Computing, Vienna, AUT) using the lme4 package (Bates & Maechler, 2015) and significance was calculated using the lmerTest package (Kuznetsova *et al.*, 2017), which applies Satterthwaite’s method to estimate degrees of freedom and generate p-values for mixed effects models by comparing the full model including the effect of interest against a null model excluding the effect of interest.

We used linear mixed effects models to determine if age group (categorical) and muscle were significant predictors for our MU variables. We employed age group (younger vs. older), muscle (BIC vs TRI), and their interaction as fixed effects. As random effects, we included a random intercept for each subject as well as a random slope accounting for the muscle within each subject. Effects estimated from the linear mixed effects models are presented as parameter estimates ± SE.

In order to avoid the bimodal distribution of ages created by our selective sampling of healthy younger and older adults, we used separate generalized linear mixed effects models (computed by the MuMIn R package (Barton, 2018) to identify significant relationships between ΔF and age in the younger and older groups. This was done to assess the degree to which the age of the subjects in each group could account for variance in ΔF. Specifically, we analyzed whether age was able to predict ΔF. For each age group, we included age (continuous variable), muscle (TRI vs. BIC), and their interaction as fixed effects. As random effects, we included a random intercept for each subject as well as a random slope accounting for the muscle within each subject. Variance accounted for by the model is reported as conditional R^2^GLMM values, whereas variance accounted for by only the fixed effects is reported as marginal R^2^GLMM values (Nakagawa & Schielzeth, 2013; Johnson, 2014; Nakagawa & Schielzeth, 2017).

## RESULTS

In the ten younger participants, decomposition yielded 1002 MU spike trains from the BIC, and 1211 MU spike trains from the TRI. In the ten older participants, decomposition yielded 533 MU spike trains from the BIC, and 827 from the TRI. All participants completed a minimum of 10 submaximal torque ramps in the directions of EF and EE. An average of 6.2 (3.81) and 9.0 (4.99) MUs per trial were decomposed from the BIC and TRI, respectively, of younger participants, and 4.4 (1.60) and 6.7 (3.39) MUs per trial were decomposed from the BIC and TRI, respectively, of older participants. Following the visual inspection and removal of erroneous spike times, the mean silhouette values of the decomposed motor units were 0.91 (0.05) from the BIC and 0.90 (0.05) from the TRI of the younger participants. The mean silhouette values from the older participants was 0.91 (0.04) from the BIC and 0.93 (0.05) from the TRI. An example of smoothed MU firing rate patterns of decomposed TRI MUs in a single trial from one younger and older individual are shown in figure 3, which shows many of the features that will be quantified below.

**Figure 3:**
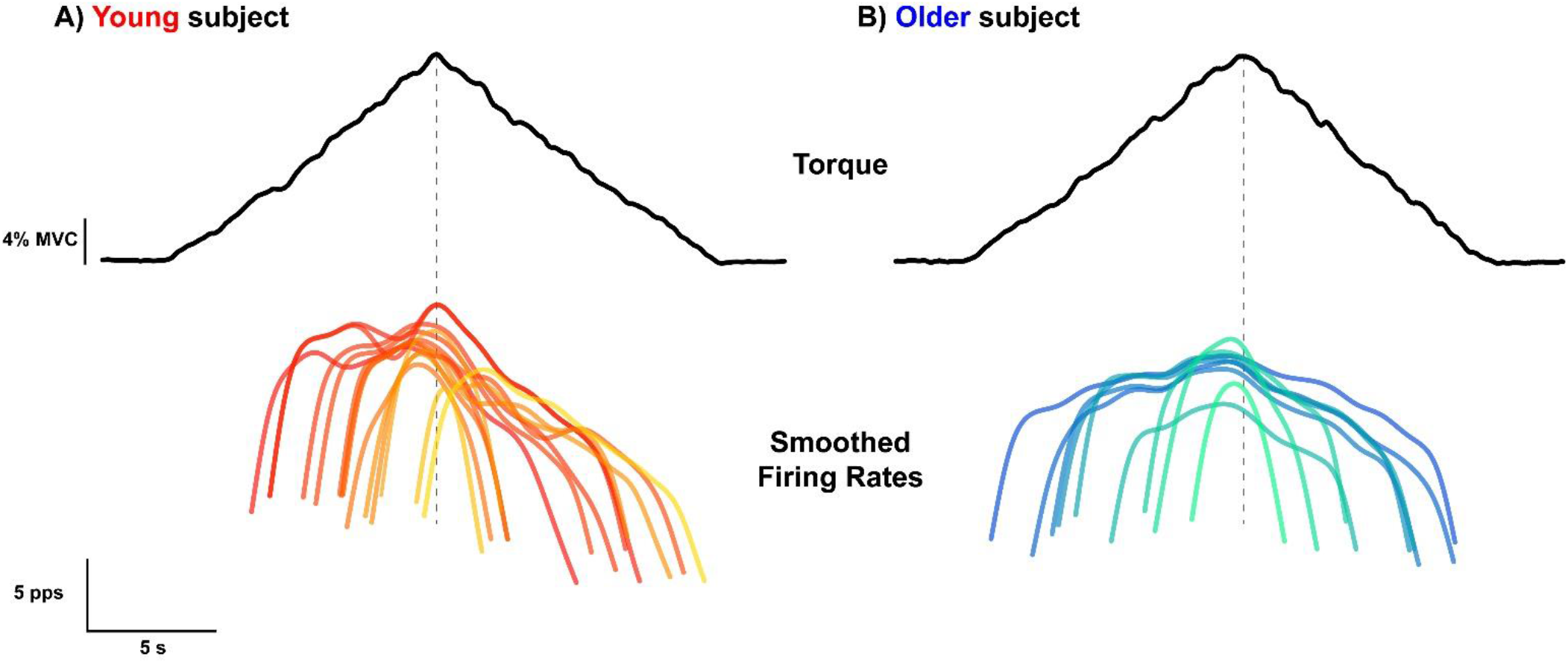
Single trial elbow extension torque (top; black traces) and smoothed firing rates of all decomposed triceps brachii MUs (bottom; coloured traces) for a younger (left; A) and an older (right; B) participant. Smoothed firing rates are darker at lower thresholds, and lighter for higher thresholds for both participants.

### Comparison of peak MU firing rates

The peak firing rates from BIC and TRI MUs are shown in Figure 4. In this, and in the following figures, each data point represents the mean value from all MUs collected from one subject during the submaximal EF torque ramps (BIC data) or submaximal EE torque ramps (TRI data). In congruence with previous findings (Dalton *et al.*, 2010), older participants had reduced peak firing rates compared to younger participants. Group mean peak firing rates for the younger participants were higher than observed in the older participants in both the BIC (16.0 [1.74] pps vs 14.5 [2.22] pps, *ES = 0.71*) and the TRI (17.9 [1.52] pps vs 16.3 [1.90] pps, *ES = 0.84*). A linear mixed effects model revealed that both age group (*χ*^*2*^ *[1] = 4.7564, P = 0.0292*) and muscle (*χ*^*2*^ *[1] = 15.731, P < 0.0001*) were significant predictors of peak firing rate, but the interaction between the two variables was not significant (*P = 0.9980*). Peak MU firing rates were lowered by 1.6 (0.72) pps (*P = 0.0412*) in older participants, and were 1.8 (0.38) pps (*P = 0.0001*) higher in TRI, compared to BIC.

**Figure 4:**
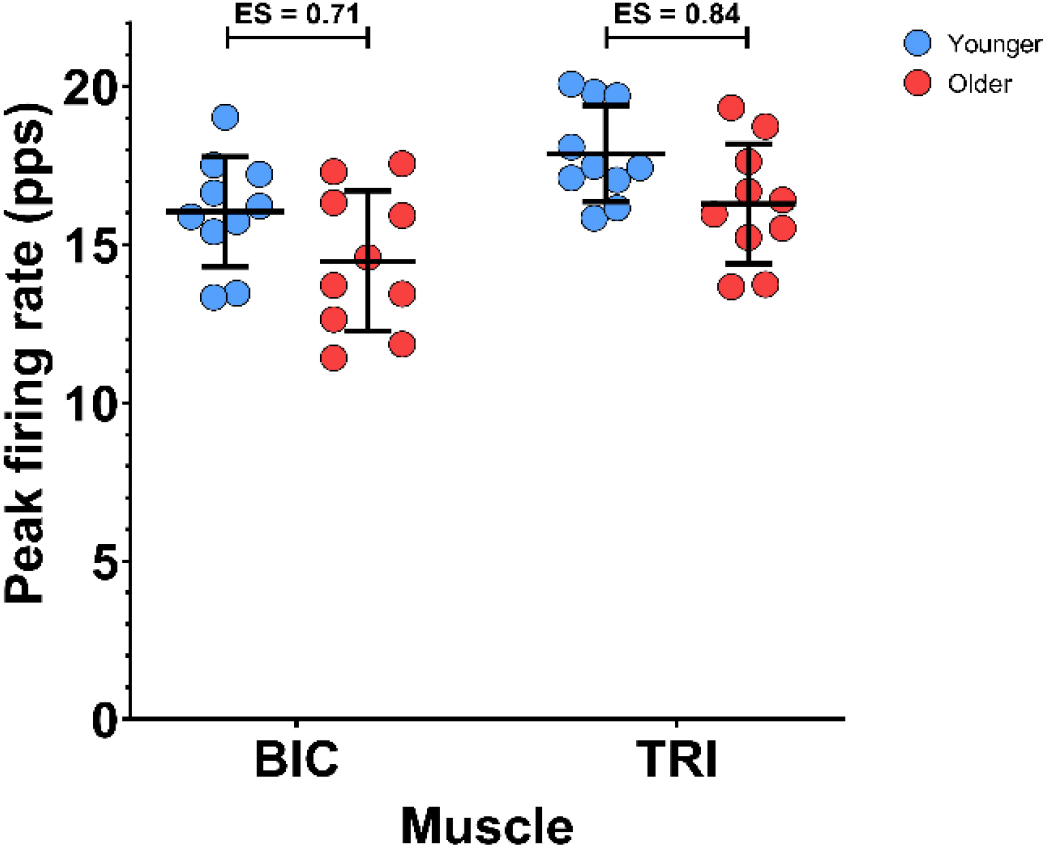
Individual participant means (younger = blue; older = red) and group mean and SD (black) for the peak firing rate of BIC (biceps brachii) MUs during a 20% elbow flexion ramp, and TRI (triceps brachii) MUs during a 20% elbow extension ramp.

### Range of MU firing rates

Figure 5A shows the range of MU firing rates from BIC and TRI MUs. The group mean range of firing rates from the younger participants were higher than the MU firing rate ranges in the older participants in both BIC (12.0 [1.45] pps vs 9.4 [1.66] pps, *ES = 1.48*) and TRI (13.6 [1.22] pps vs 10.5 [1.97] pps, *ES = 1.70*). Age group (*χ*^*2*^ *[1] = 7.7509, P = 0.0053*) and muscle (*χ*^*2*^ *[1] = 15.570, P < 0.0001*) were both significant predictors of firing rate range; the interaction of age group and muscle was not significant (*P = 0.8687*). The range of MU firing rates was 1.7 (0.60) pps (*P = 0.0091*) higher in younger participants than older participants, and 1.5 (0.32) pps (*P = 0.0001*) higher in MUs from the TRI than the BIC.

**Figure 5:**
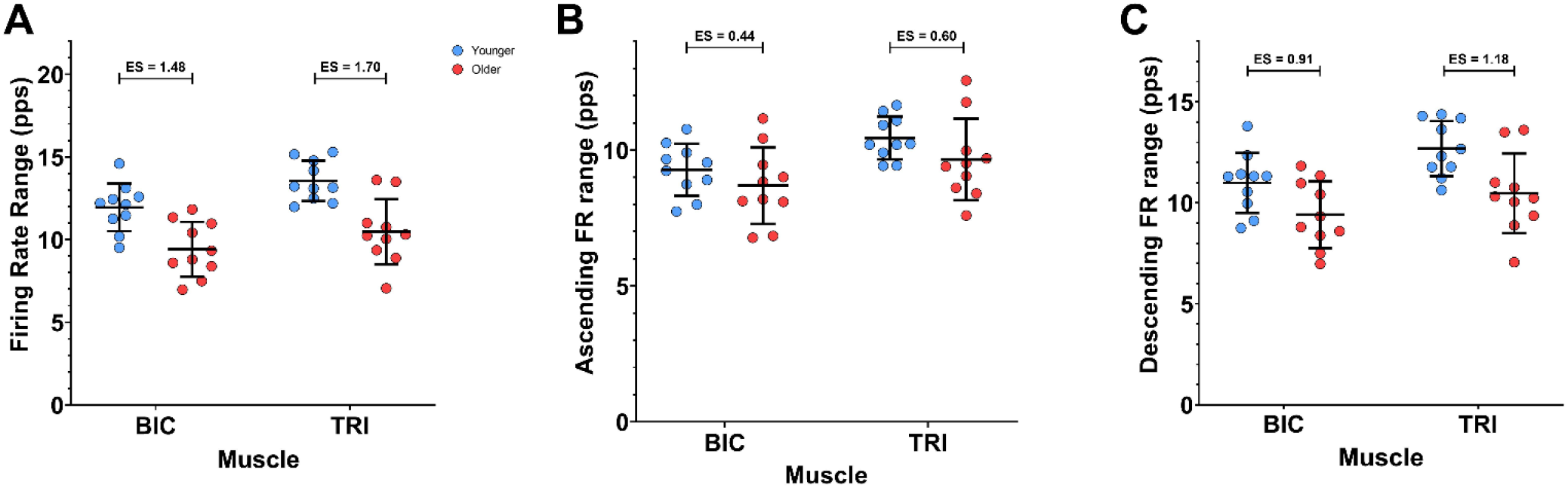
Participant means (younger = blue; older = red), along with group mean and SD (black), for the range of MU firing rates during the full ramp (A), the ascending limb of the ramp (B), and the descending limb of the ramp (C). BIC (biceps brachii) data is from elbow flexion ramps, and TRI (triceps brachii) data is from elbow extension ramps.

The observed range of MU firing rates for the ascending and descending limbs of the torque ramps are displayed in Figure 5B and 5C, respectively. During the ascending torque phase, the group mean firing rate range in the younger participants was 9.3 (0.96) pps in the BIC and 10.4 (0.79) pps in the TRI. While in the older participants, the group mean firing rate range was 8.7 (1.41) pps in the BIC and 9.7 (1.50) pps in the TRI. In the BIC, the effect size of the difference in firing rate range on the ascending limb between younger and older participants was *0.44*, and in the TRI the effect size was *0.60*. However, age group was not a significant predictor (*P = 0.1112*) nor was the interaction between muscle and age group (*P = 0.7546*). Muscle was the only significant predictor of MU firing rate range over the ascending limb of the torque ramp (*χ*^*2*^ *[1] = 10.9590, P = 0.0009*). The ascending limb firing rate range was 1.1 (0.29) pps higher in the TRI than in the BIC (*P = 0.0014*).

During the descending torque phase, the group mean MU firing rate ranges were higher in the younger participants compared to the older participants for both muscles (BIC: 11.0 [1.49] pps vs 9.4 [1.66] pps, *ES = 0.91*; TRI: 12.7 [1.37] pps vs 10.5 [1.97] pps, *ES = 1.18*). Both age group (*χ*^*2*^ *[1] = 7.7199, P = 0.0055*) and muscle (*χ*^*2*^ *[1] = 10.856, P = 0.0010*) were significant predictors of descending limb firing rate range, in our model. The interaction between age group and muscle was not significant (*P = 0.4002*). During the descending torque ramps, the firing rate range was 1.9 (0.62) pps higher in younger participants (*P = 0.0083*), as compared to older participants, and 1.4 (0.37) pps higher in the TRI (*P = 0.0015*), as compared to the BIC.

Similar to the peak firing rates, older participants showed a reduced range of MU firing rates overall, as well as a reduction in firing rate range on the descending limb. However, the firing rate range during the ascending portion of the torque ramp was not significantly affected by aging. The difference in firing rate range between younger and older participants can be appreciated in an example of smoothed firing rates from the TRI of one younger and one older participant in figure R1.

### Estimates of PIC amplitude using ΔF

Subject mean values for the ΔF calculation are shown in Figure 6. Group means for ΔF were substantially higher in the younger participants than in the older participants in the BIC (4.1 [1.35] pps vs 2.3 [0.84] pps, *ES = 1.47*) and in the TRI (5.2 [0.94] pps vs 3.2 [1.10] pps, *ES = 1.84*). Age group (*χ*^*2*^ *[1] = 18.326, P < 0.0001*) and muscle (*χ*^*2*^ *[1] = 17.796, P < 0.0001*) were both significant predictors for ΔF, in our model, however the interaction between those variables was not significant (*P = 0.2848*). ΔF was reduced by 1.9 (0.36) pps in the older participants (*P < 0.0001*), and 1.3 (0.24) pps lower in the BIC than in the TRI (*P < 0.0001*).

**Figure 6:**
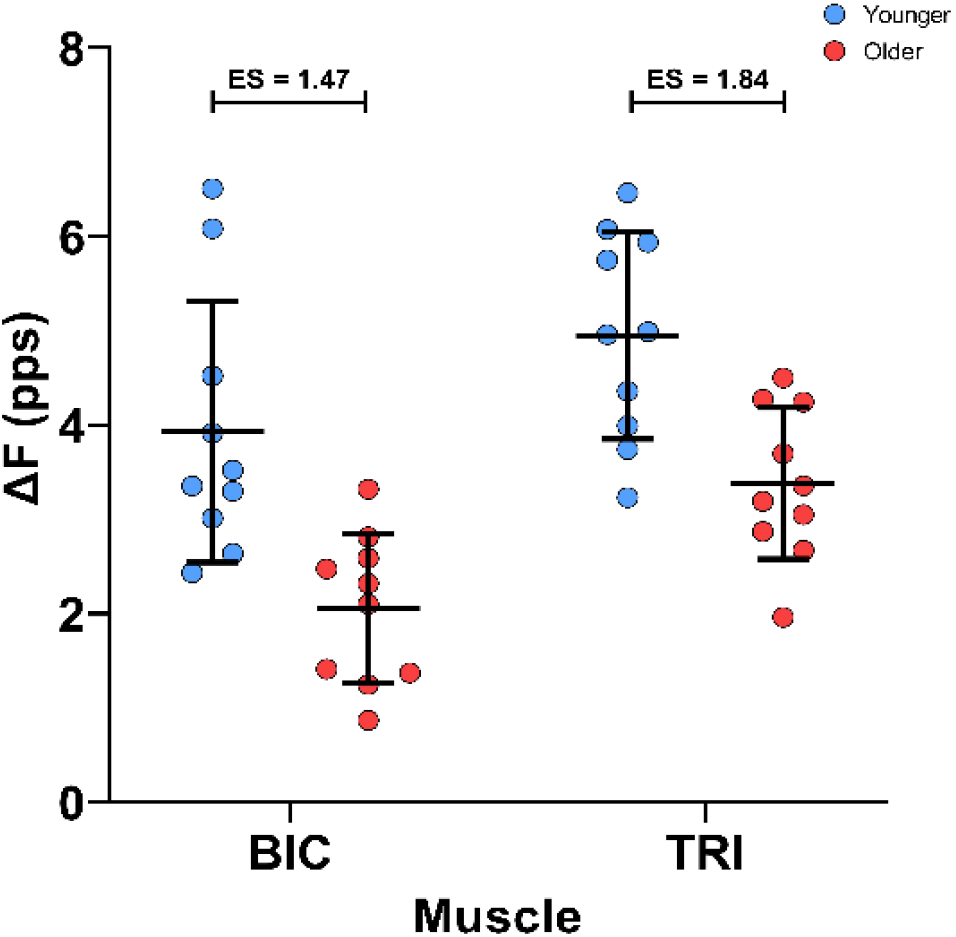
ΔF estimates from the BIC (biceps brachii) during elbow flexion and TRI (triceps brachii) during elbow extension. Participant means in color (younger = blue; older = red), with the black bars denoting group mean and SD.

We then determined whether any relationship existed between the reported age of participants and ΔF within each of these age groups. In Figure 7, ΔF is plotted as a function of participant age along with regression lines from our model. In the younger participants, the generalized linear mixed effects model accounted for 31.96% of the variance in ΔF, with the fixed effects of muscle and age accounting for 5.28% of the variance. Muscle was a significant predictor of ΔF (*χ*^*2*^ *[1] = 5.3981, P = 0.0202*), however, age was not a significant predictor of ΔF in the younger participants (*χ*^*2*^ *[1] = 0.0021, P = 0.9637*). The interaction between age and muscle was also not a significant predictor (*χ*^*2*^ *[1] = 0.3742, P = 0.5407*).

**Figure 7:**
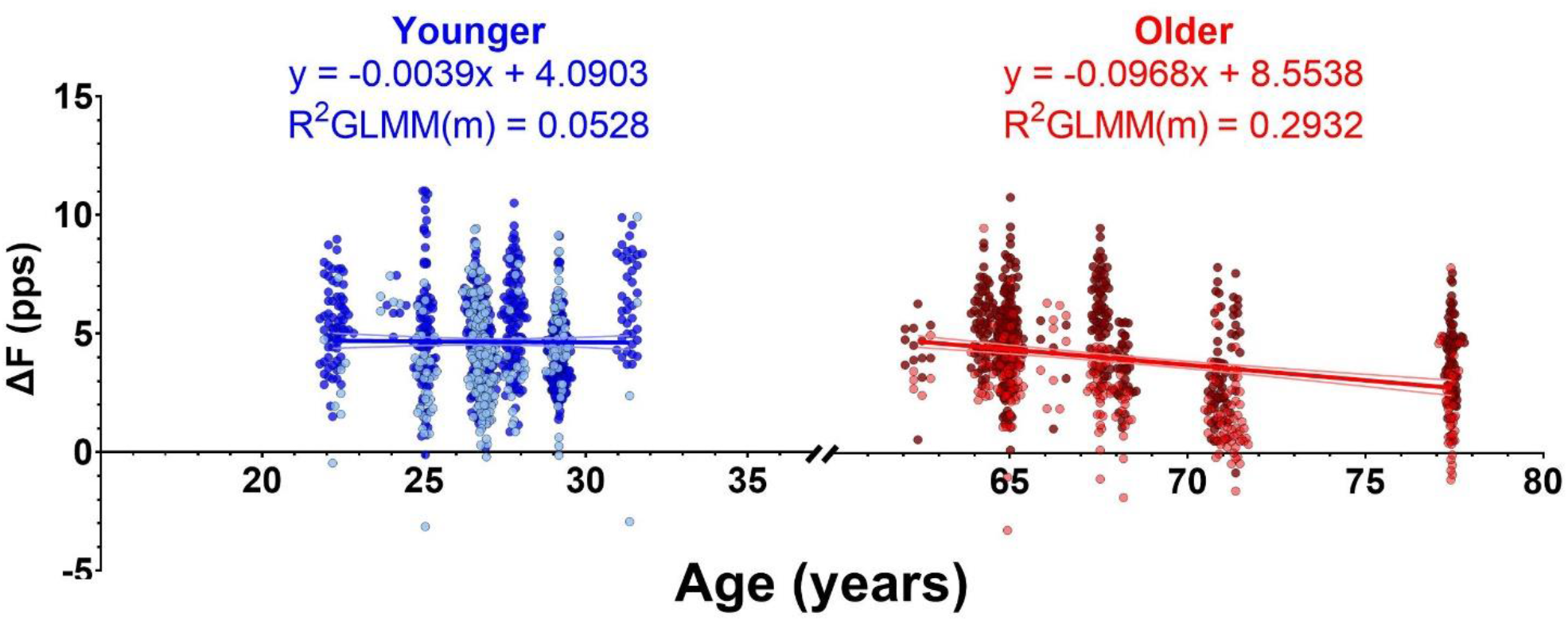
The relationship between ΔF and age of the younger (blue; left) and older (red; right) group participants. The lines indicate individual generalized linear mixed effects models. We show only the overall slope of the model, which includes fixed effects of age and muscle. For clarity of display, individual participants are not distinguished from one another, however the random effect in the model does account for the variability within each participant. Dark and light data points indicate ΔF values from TRI (triceps brachii) and BIC (biceps brachii) MUs, respectively. Some jitter was added to the data points x-value (age) for clarity of display Equations derived from the model are displayed for the younger and older participants along with the marginal R^2^GLMM, which indicates the variance accounted for by our fixed effects of muscle and age.

In the older participants, the model accounted for 45.98% of the observed variance in ΔF, with the fixed effects accounting for 29.32% of the variance. Both muscle (*χ*^*2*^ *[1] = 14.75, P = 0.0001*) and age (*χ*^*2*^ *[1] = 4.9504, P = 0.0261*) were significant predictors of ΔF, but the interaction between them was not (*χ*^*2*^ *[1] = 0.8031, P = 0.3702*). Greater reductions in ΔF were associated with increasing age, in the older participants (−0.097 [0.041] pps/year, *P = 0.0473*). In summary, ΔF was reduced in older participants compared to younger participants and a negative relationship existed between age and ΔF in the older participants, but not in the younger participants.

### MU firing duration

Provoked by the reduction in firing rate hysteresis in older participants (i.e. reduced ΔF), we investigated whether the duration of the MU firing differed between age groups; the subject and group means for MU duration are shown in Figure 8A. The group means for total MU firing duration in the younger participants was 9.7 (1.95) s in the BIC and 8.9 (1.38) s in the TRI. In the older participants, the group mean for MU firing duration was 9.8 (1.83) s in the BIC and 9.6 (1.64) s in the TRI. The effect sizes between younger and older participants are *0.05* for the BIC and *0.43* in the TRI. However, the linear mixed effects model found that age group (*P = 0.3534*), muscle (*P = 0.3182*), and the interaction between age group and muscle (*P = 0.5084*) were not significant predictors of MU firing duration.

**Figure 8:**
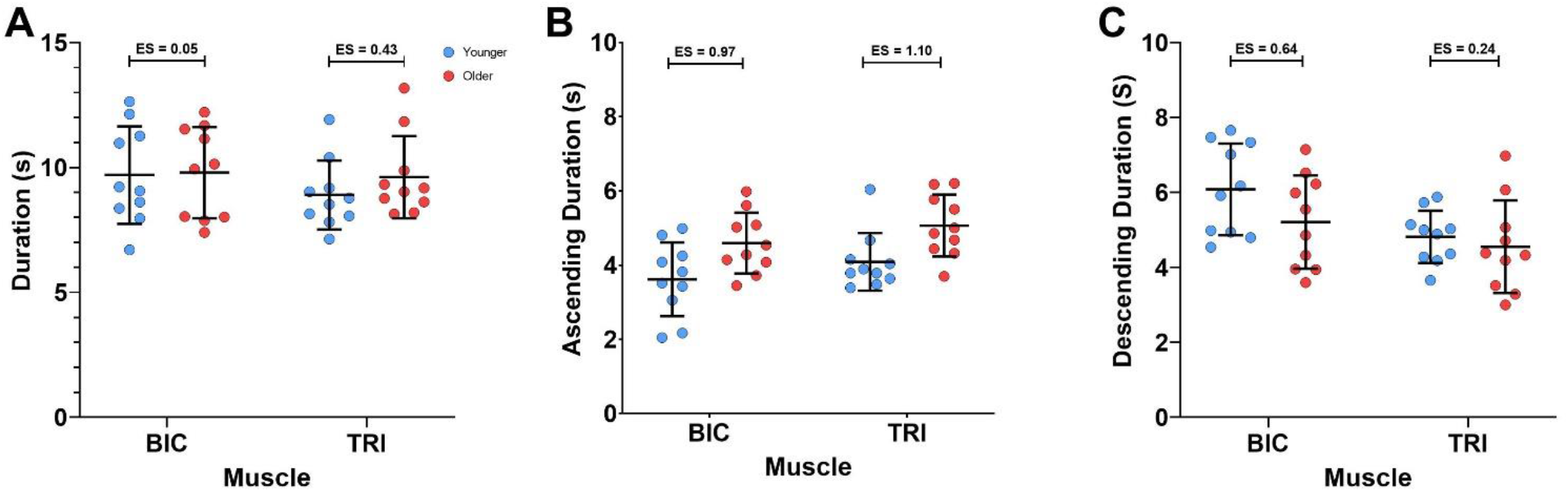
Participant means (color) and group means (black), showing the firing duration of MUs for the full torque ramp (A), the ascending limb of the torque ramp (B), and the descending limb of the torque ramp (C).

MU firing was further investigated by comparing the duration of firing during the ascending and descending limbs of the torque ramps, as shown in Figure 8B and C, respectively. On the ascending limb of the ramp, the group means for MU firing duration were shorter in the younger participants than in the older participants (see Figure 3 for example) in both the BIC (3.6 [0.99] s vs 4.6 [0.82] s, *ES = 0.97*) and in the TRI (4.1 [0.78] s vs 5.1 [0.83] s, *ES = 1.10*). Our model found age group (*χ*^*2*^ *[1] = 9.3990, P = 0.0022*) and muscle (*χ*^*2*^ *[1] = 4.5555, P = 0.0328*) were both significant predictors of firing duration on the ascending limb of the torque ramp, while the interaction between the two variables was not significant (*P = 0.9224*). Ascending limb firing duration was increased by 1.0 (0.30) s in the older participants (*P = 0.0039*), compared to younger participants, and 0.5 (0.22) s longer in TRI MUs than BIC MUs (*P = 0.0401*).

On the descending limb of the torque ramps, the group means for MU firing duration were longer for the younger participants than the older participants in both muscles (BIC: 6.1 [1.22] s vs 5.2 [1.24] s, *ES = 0.64*; TRI: 4.8 [0.70] s vs 4.6 [1.24] s, *ES = 0.24*). Muscle was revealed to be a significant predictor of firing duration on the decreasing torque ramp (*χ*^*2*^ *[1] = 7.4705, P = 0.0063*), but age group was not a significant predictor (*P = 0.1765*) and the interaction of age group and muscle was not significant (*P = 0.3589*). Descending limb firing duration was 1.0 (0.33) s longer in BIC MUs than in TRI MUs.

Age was only associated with an increased duration of firing on the ascending limb of the torque ramp, but did not significantly affect the overall firing duration or the firing duration on the descending limb of the torque ramp. As shown in figure 3, a longer duration of firing during the ascending phase of the ramp without a difference during the descending phase of the ramp or total duration of firing would indicate a leftward shift, and more symmetrical pattern of firing.

## DISCUSSION

The present study aimed to investigate the effects of healthy aging on the MU firing patterns from the biceps brachii (BIC) and triceps brachii (TRI) by comparing younger and older healthy adults. In agreement with previous literature, we have found lower peak firing rates in both the BIC and TRI of older adults. Further, and perhaps most novel, we found substantial and significant reductions in estimates of persistent inward currents (PICs; ΔF) in older adults irrespective of muscle. In older adults, we also found that age was a significant predictor of ΔF (i.e. ΔF decreased with respect to age); however, there was no such relationship in the younger group. Additional characteristics of MU firing patterns support the notion of reduced onset-offset hysteresis, such as reduced firing rate range during the descending phase of the ramp and a leftward shift in MU firing in older compared to younger adults. These findings suggest that MU firing patterns in older people exhibit less PIC activity, which may have implications for motor control.

### Reduced motor unit firing rates in older people

Reductions in MU firing rates in older adults here are similar to those reported in previous studies. For example, using intramuscular EMG, Dalton et al (Dalton *et al.*, 2010) found that mean firing rates are reduced in both the BIC and TRI across a variety of voluntary effort levels. The difference between younger and older adults was substantial (~45%) at high levels of effort for both muscles, but the differences at 25% MVC were modest and similar in magnitude to what we observed. Indeed, firing rates have been shown to be reduced in older adults in other studies as well, with larger differences at higher contraction intensities (Kamen *et al.*, 1995; Connelly *et al.*, 1999; Kamen & Knight, 2004; Barry *et al.*, 2007; Kirk *et al.*, 2018; Kirk *et al.*, 2019).

Organizational changes to the MUs following age-related loss of motoneurons (Doherty *et al.*, 1993) could play a role in the observed reductions in firing rate. With re-modelling there are changes in MU twitch contraction times such that twitch fusion occurs at a lower rate (Bellemare *et al.*, 1983; Newton *et al.*, 1988; Connelly *et al.*, 1999; Roos *et al.*, 1999). However, investigations into motoneuron loss with age have shown a very moderate loss in individuals below the age of 60 (Tomlinson & Irving, 1977). As the mean age of the older participants in this group is only 67.8, some of our sample may not have substantial loss of motoneurons, and the subsequent reorganization of the motor pool would lead to only modest reductions of firing rates.

Changes in the pattern or level of excitatory synaptic input to the motoneurons are also likely to contribute to the observed reduction in firing rates with age. For instance, excitatory post-synaptic potential (i.e. EPSP) amplitudes, as estimated with peristimulus time histograms, in response to transcranial magnetic stimulation of the motor cortex, are reduced by ~50% in older people (> 55 years old). Hypoexcitability of the corticospinal pathway corresponds with weakness in older adults (Clark *et al.*, 2015), and ~33% of the variance in weakness in seniors can be accounted for by measures of corticospinal excitability (i.e. motor evoked potential amplitudes) and inhibition (i.e. corticospinal silent period). Stretch reflexes are also reduced with aging, suggesting that homonymous 1a input from muscle spindles is reduced (Bryndum & Marquardsen, 1964; Milne & Williamson, 1972), and slowed (i.e. longer half relaxation times) with advancing age (Carel *et al.*, 1979), which could be due to a combination of changes in muscle spindle sensitivity (Swash & Fox, 1972), Ia-motoneuronal transmission, and/or spinal motoneuron excitability (Geertsen *et al.*, 2017). Interestingly, the Hoffman (H-) reflex, which bypasses the muscle spindles by direct peripheral nerve stimulation, is also reduced with age, providing further support for Ia-motoneuronal transmission and/or spinal motoneuron excitability alterations with age (deVries *et al.*, 1985; Scaglioni *et al.*, 2003). Reductions in any combination of the above mentioned excitatory input would surely result in reduced MU firing rates in older adults.

Inhibitory spinal circuits can also affect descending input and/or Ia afferent input to spinal motoneurons via inhibitory interneurons. Older adults also have reduced indices of reciprocal inhibition, both at the spinal (Kido *et al.*, 2004) and cortical levels (Hortobagyi *et al.*, 2006), which may alter the commands to agonist-antagonist muscle pairs. Although the effects of aging on Ia presynaptic inhibition at rest are unclear (Butchart *et al.*, 1993; Morita *et al.*, 1995; Earles *et al.*, 2001), there is a decrease in the amount of modulation of Ia presynaptic inhibition with increasing contraction intensity in older adults (Butchart *et al.*, 1993), which would lead to differences in the pattern of inhibition and have profound effects on the balance of excitation, inhibition, and neuromodulation (Johnson *et al.*, 2017) required to perform a task. The imbalance of inhibition and excitation could therefore play a role in the reduced firing rates we observed in our older adults.

MU firing rates are highly dependent upon the biophysical properties of the parent motoneurons, and age-related changes of such properties can lead to reductions in firing rates. Spike after-hyperpolarization (AHP) duration is increased in aged rodents (Cameron *et al.*, 1991; Kalmar *et al.*, 2009) and cats (Morales *et al.*, 1987). Similarly, AHP duration increases gradually with age (Piotrkiewicz *et al.*, 2007), and when compared directly, AHP is longer in older compared to younger adults (Christie & Kamen, 2010), which can contribute to reductions in MU firing rates. Other biophysical properties that are altered with age can affect recruitment and repetitive firing of MUs, such as increases in the input resistance (Chase *et al.*, 1985; Morales *et al.*, 1987; Kalmar *et al.*, 2009), and reduced rheobase current (Morales *et al.*, 1987; Kalmar *et al.*, 2009), suggesting that aged motoneurons are less excitable. Indeed motoneuron recruitment gain is reduced with age, as evidenced in a recent investigation by Nielsen and colleagues (Nielsen *et al.*, 2019). Therefore, changes to the biophysical properties of the motoneurons likely contribute to the reduced firing rates we observed in older adults.

Not only were peak firing rates reduced in the current investigation, but older adults also showed a compressed range of MU firing rates. This compressed range of firing can arise from similar mechanisms that underlie the reductions in peak firing rates, but it is interesting to note that the firing rate range during the ascending phase of the contraction was similar for both younger and older adults. That is, the rate modulation from the onset of firing to peak firing was similar (see figure 5B). On the contrary, the reduction in overall firing rate range (~1.7 pps) seen in older participants, is primarily attributed to the reduction in firing rate range seen on the descending limb of the torque ramp (~1.9 pps). The reduced firing range seen on the descending limb (I.e. reduced hysteresis) of the torque ramps is most likely related to a decrease in PIC activity, which brings us to the next topic of our discussion (i.e. reduced estimates of PICs in older adults).

### Reduced estimates of PICs in older people

We have shown that ΔF is substantially lower, regardless of muscle, in the upper limb of older adults, compared to younger adults. Further, in our older group, we found that increasing age was associated with reductions in ΔF (see figure 7). In addition, the relative firing duration of motor units is shifted to the left in older adults, such that the duration of firing is symmetrical during the ascending and descending phases of the ramp contraction, indicating less onset-offset hysteresis. Although ΔF and the leftward shift in firing patterns are an indirect estimate of PIC activity, they do support our hypothesis that PICs are reduced older people. Such age-related changes in estimates of PICs are most likely due to either changes in 1) monoaminergic input to the motoneurons, 2) the amount or pattern of inhibition, and/or 3) Na+ or Ca2+ channel function.

The overall leftward shift in the firing patterns of older individuals provides further insights into the effects of aging on MU firing. Based on the reduced ΔF and reduced firing rate range on the descending limb in the older adults, we expected to see a reduced duration of firing on the descending limb in those participants. Instead, we found an increased duration of firing on the ascending limb of the torque ramps in older participants, without significant changes to the overall duration and descending duration of firing, which may lead to the observed ΔF differences. This may suggest that motor units in older participants were recruited earlier than those from younger participants. Indeed, this would be similar to other reports that have shown a lower average recruitment threshold for motor units recorded from older compared to younger adults (Erim *et al.*, 1999; Klass *et al.*, 2005, 2008; Fling *et al.*, 2009; Pascoe *et al.*, 2011). Most intriguing though is the fact that the firing patterns of older individuals were more symmetrical (~approximately equal time on the ascending and descending portion of the ramps – see figure 3), indicating less hysteresis, a behaviour that is a hallmark of PICs.

Schwindt and Crill (Schwindt & Crill, 1980) first discovered PICs in motoneurons by blocking K^+^ currents in anesthetized preparations, but by using un-anesthesthetized decerebrate preparation, Hounsgaard and colleagues (Hounsgaard *et al.*, 1984; Hounsgaard & Kiehn, 1989) showed that PICs emerged in the presence of serotonin (5HT) or norepinephrine (NE), meaning that PICs are a natural consequence of endogenous neurotransmitters. Although important for initiation of repetitive firing, the component of the PIC mediated by Na+ is not responsible for the long-lasting prolongation of firing due to the inactivation time constant of just a few seconds (Lee & Heckman, 1998; Binder & Powers, 2001). Instead, it is the component mediated by L-type Ca2+ channels that causes prolonged self-sustained firing and profound onset-offset hysteresis (Lee & Heckman, 2001; Harvey *et al.*, 2006; Kuo *et al.*, 2006), in particular, the Ca2+V1.3 channels show little or no inactivation (Moritz *et al.*, 2007; Binder *et al.*, 2020). This “self-sustained” firing, or simply maintained motoneuron firing with decreased synaptic input, was emphasized as being quite important for postural behaviours because sustained force generation can be maintained without the need for sustained descending input (Hounsgaard *et al.*, 1984; Hounsgaard & Kiehn, 1989). Therefore, the functional role of PICs in normal human behaviours can easily be appreciated (Heckman *et al.*, 2008b). With a reduction in PIC-related prolongation of motoneuron firing, simple tasks would require greater synaptic input to maintain activities such as standing or carrying objects. Since the aging process is associated with reduced excitatory input to motoneurons, the role of PICs in maintaining forces required for everyday activities may be increased with advancing age.

Initial attempts to understand PIC effects of human MU firing behaviour focused on self-sustained firing or “bi-stability”(Kiehn & Eken, 1997). In such experiments, MUs are tracked during low-level voluntary efforts and an additional source of synaptic input (i.e. vibration) causes the recruitment of an additional MU (test unit) that maintains firing after the additional input is removed (Gorassini *et al.*, 1998). MUs can then be classified as either having PICs, or not, based on the occurrence of test units that maintain firing after the additional synaptic input is removed. Using this approach, Kamen and colleagues (Kamen *et al.*, 2006) showed that older individuals have a similar occurrence (23.1%) of MUs that exhibit self-sustained firing as younger adults (22.8%). As such, and contrary to the findings in our current investigation, they concluded that PIC-like behaviour does not seem to be affected by the aging process. They did, however, report that the mean drop-out torque of newly recruited MUs was slightly higher for older adults (3.26% maximal voluntary contraction (MVC) vs 2.43% MVC), although variability was high and therefore no statistical differences were reported. It is important to note that the occurrence of self-sustained firing may not be the be-all-end-all method to quantify whether PICs are present during voluntary motor behaviour in humans. This is because PICs almost certainly contribute to motoneuron firing during all voluntary behaviours, because without the amplification effects of PICs, the small currents produced by descending and sensory inputs are too weak to have much of an effect on motonneuron firing (Binder & Powers, 2001). More important to the understanding of human motor output is the magnitude of PICs, rather than the presence.

Hysteresis of MU firing rates, on the other hand, has proven to be the most consistent hallmark for non-invasive estimation of the magnitude of PICs in humans, as was first realized by Gorassini, Bennett and colleagues (Bennett *et al.*, 2001a; Gorassini *et al.*, 2002). The now standard paired-MU analysis technique (ΔF) has been subject to rigorous investigations interested in the accuracy of these estimates (Bennett *et al.*, 2001a; Bennett *et al.*, 2001b; Powers *et al.*, 2008; Revill & Fuglevand, 2011; Powers & Heckman, 2015; Afsharipour *et al.*, 2020; Hassan *et al.*, 2020). Bennett and colleagues (Bennett *et al.*, 2001a; Bennett *et al.*, 2001b) used parallel MU and intracellular recordings in rat motoneurons to clearly demonstrate that ΔF reflects features of PICs. With advances in technology, these estimates of PICs have been obtained across hundreds of MUs (Afsharipour *et al.*, 2020; Hassan *et al.*, 2020; Kim *et al.*, 2020; Trajano *et al.*, 2020), which likely provides a better overall estimation of PIC magnitude across the entire motor pool. Even though MUs of older adults in our experiment certainly displayed onset-offset hysteresis (i.e. positive ΔF values overall), the magnitude of this hysteresis was markedly reduced compared to the sample of younger adults that were recruited. In fact, estimates of PICs (ΔF) were reduced by ~40%, with very large effect sizes (all ES > 1.45). In addition, the age of individuals was a significant predictor of ΔF, suggesting that PICs may deteriorate with age at a rate of ~1pps/decade, but only in older adults.

The magnitude of PICs is directly proportional to the level of NE and 5HT (Lee & Heckman, 1998, 2000), which are monoamines released from the from the caudal raphe nucleus and locus coeruleus, respectively. These monoaminergic nuclei of the brainstem deteriorate with age (Shibata *et al.*, 2006; Pagano *et al.*, 2017), and in particular, the age-related reduction in locus coeruleus structural integrity is associated with impaired cognitive and behavioural function (Liu *et al.*, 2020), as well as reductions in central pain modulation (Grashorn *et al.*, 2013; Damien *et al.*, 2018). Deterioration of these nuclei could also lead to reductions in neuromodulatory drive to motorneurons, reducing PIC activity, which would ultimately explain some of the reductions observed in ΔF. NE-mediated effects are likely predominantly due to degradation of the locus coeruleus because older rodents maintain only ~30% NE nuclei compared to ~90% 5HT nuclei (Tatton *et al.*, 1991). Despite the evidence that a greater proportion of raphe nuclei are maintained with age, spinal 5HT is greatly reduced (Johnson *et al.*, 1993; Ko *et al.*, 1997). Therefore, the potential of 5HT-mediated effects with the aging process are more likely to occur peripherally. With aging, there is increased circulation of cytokines (so-called “inflamm-aging”) (Michaud *et al.*, 2013), which affect 5HT receptors and increase re-uptake of 5HT (Steinbusch *et al.*, 2021). In sum, less availability of monoamines would result in reduced PIC magnitude at the same relative effort, which is what we observed as a reduction in ΔF.

PICs are also highly sensitive to inhibitory inputs (Hultborn *et al.*, 2003; Kuo *et al.*, 2003; Heckman *et al.*, 2008a; Hyngstrom *et al.*, 2008; Revill & Fuglevand, 2017). Thus, changes to the amount or pattern of inhibition may lead to reduced estimates of PICs as estimated by ΔF. As mentioned above, there are age-related alterations in spinal and supraspinal inhibitory circuits (Butchart *et al.*, 1993; Kido *et al.*, 2004; Hortobagyi *et al.*, 2006) that could modify the synaptic input to motoneurons. While difficult to measure, the temporal pattern of inhibitory commands can also affect the ΔF estimate (Powers *et al.*, 2012; Johnson *et al.*, 2017). Push-pull inhibition, where inhibition varies inversely with excitation, can lead to reductions in MU hysteresis (Powers *et al.*, 2012). It, therefore, remains possible that the pattern of the inhibitory commands is altered with age to compensate for the various structural and functional changes in the neuromuscular system (Hepple & Rice, 2016; McNeil & Rice, 2018) associated with the aging process and may contribute to our observed reductions in ΔF.

Alterations in the integrity and function of 5HT/NE receptors and voltage sensitive ion channels must also be considered in relation to age-related changes in the nervous system. Basic (i.e. larger and longer AHP, lower rheobase, greater input resistance) and rhythmic (i.e. slower minimum and maximum steady-state firing frequencies and lower f-I slopes) motoneuron properties are consistent with reduced motoneuron excitability in very old (>30 months) rodents. However, Kalmar et al (Kalmar *et al.*, 2009) also showed an increased incidence of PIC-like behaviour in very old rodent motoneurons, which they suggested to have resulted from increased 5HT and NE receptor sensitivity to residual endogenous monoamines (Harvey *et al.*, 2006) as a compensatory mechanism to counteract the reduced motoneuron excitability. Although this increased incidence of PIC may seem to contradict our findings, this type of analysis simply determines the relative number of motoneurons that have hysteresis in response to current injection to the soma, whereas we quantified the average magnitude of hysteresis during voluntary activation (i.e. axo-dendritic synaptic input). L-type Ca2+ channels are concentrated in the dendritic tree, far away from the soma, meaning that the levels of injected current may have underestimated PICs in a healthy younger motoneuron receptor hypersensitivity due to the inability to activate PICs from the soma (Bennett *et al.*, 1998; Lee & Heckman, 1998). Aging may also result in changes in the expression of receptor subtypes or the downstream signaling of various receptors. Indeed, there are age-related reductions in the duration of Ca2+-mediated plateau potentials in striatal neurons (Dunia *et al.*, 1996), and more generally, deregulated Ca2+ is an active component of healthy aging that can increase the risk of cell death and neurodegenerative disorders (Nikoletopoulou & Tavernarakis, 2012). As such, it is difficult to pinpoint the exact monoaminergic receptor or ion channel dysfunction that may contribute to the observed reductions in estimated PICs during voluntary contractions with aging.

### Methodological considerations

Since this experiment was conducted in a non-invasive fashion, we were unable to directly determine the PIC magnitude. Instead, we relied on the best estimation of PICs available in humans (i.e. ΔF), which has undergone rigorous scrutiny to ensure accuracy of the estimates (Bennett *et al.*, 2001a; Bennett *et al.*, 2001b; Powers *et al.*, 2008; Revill & Fuglevand, 2011; Powers & Heckman, 2015; Afsharipour *et al.*, 2020; Hassan *et al.*, 2020). Nonetheless, it is difficult to determine whether the reduction in ΔF is the result of alterations in monoaminergic drive, the amount or pattern of inhibition, and/or changes to the monoaminergic receptor sensitivity or ion channel function. This delineation will require further work.

Overall, the MUs decomposed in the older participants had longer durations on the ascending phase of the ramp, suggesting a lower relative threshold of units decomposed for older adults. This relatively lower recruitment torque could lead to a ceiling effect in terms of how much hysteresis those units could exhibit. However, as the average duration of MU firing on the descending limb was 5.2 s and 4.6 s for the BIC and TRI respectively in the older participants with the descending limb of the torque ramp being 10 seconds long, we do not believe the reduced ΔF observed was due to early derecruitment due to the time constraints of the task.

### Practical considerations

In the words of Power and colleagues (Power *et al.*, 2016) – “If you don’t use it, you’ll likely lose it.” Whether this holds true for PICs is unclear at the moment, although Latella (Latella, 2021) recently made a compelling argument for studying the efficacy of strength training to mitigate the effects of aging on MU firing behaviour. Indeed, the work of Power and colleagues (Power *et al.*, 2010) suggests that estimates of MU numbers are greater in masters runners compared to their sedentary counterparts. Further, strength training-induced plasticity of motoneurons is not limited to younger adults. AHP duration is longer in older compared to younger adults, but that duration can be reduced with strength training in both age groups (Christie & Kamen, 2010). Thus, it remains possible that strength training, which necessitates high levels of effort (likely utilizing high levels of monoaminergic drive), could mitigate deterioration of monoaminergic function and/or PIC behaviour seen in older adults.

## CONCLUSION

The present study compared the firing patterns of MUs from the elbow extensors and flexors of healthy younger and older adults during isometric ramp contractions. Irrespective of muscle, age was a significant predictor of peak firing rate and firing rate hysteresis, such that both were reduced in older adults. In addition to the differences observed between age groups, the age of individuals within the older group predicted a ~1pps per decade reduction in ΔF, a non-invasive estimate of PIC magnitude across the motor pool. This reduced estimate of PIC magnitude likely arises from reductions in monoaminergic input, alterations in the amount or pattern of inhibition, and/or alterations in monoamine receptor or ion channel function. It remains unclear whether alterations in firing rate hysteresis are a compensatory adjustment or impairment that occurs with aging, however, it remains possible that physical training may be able to mitigate such changes.

